# Improving Memory via Automated Targeted Memory Reactivation during Sleep

**DOI:** 10.1101/2022.06.28.497977

**Authors:** Nathan W. Whitmore, Jasmine C. Harris, Torin Kovach, Ken A. Paller

## Abstract

A widely accepted view in memory research is that previously acquired information can be reactivated during sleep, leading to persistent memory storage. Recently, Targeted Memory Reactivation (TMR) has been developed as a technique whereby specific memories can be reactivated during sleep using a sensory stimulus linked to prior learning. TMR can improve various types of memory, raising the possibility that it may be useful for cognitive enhancement and clinical therapy. A major challenge for the expanded use of TMR is that experimenters must manually control stimulation timing and intensity, which is impractical in most settings. To address this limitation, we developed the SleepStim system for automated TMR in the home environment. SleepStim includes a smartwatch to collect movement and heart-rate data, plus a smartphone to emit auditory cues. A machine-learning model identifies periods of deep non-REM sleep and triggers TMR sounds within these periods. We tested whether this system could replicate the spatial-memory benefit of in-lab TMR. Participants learned the locations of objects on a grid, and then half of the object locations were reactivated during sleep over three nights. In an experiment with 61 participants, the TMR effect was nonsignificant but varied systematically with stimulus intensity; low-intensity but not high-intensity stimuli produced memory benefits. In a second experiment with 24 participants, we limited stimulus intensity and found that TMR reliably improved spatial memory, consistent with effects observed in laboratory studies. We conclude that SleepStim can effectively accomplish automated TMR and that avoiding sleep disruption is critical for TMR benefits.

## Introduction

Sleep has long been recognized as important for good memory function (e.g., Patrick & Gilbert, 1896), but much remains to be learned about why. A prevalent view at the present time is that reactivation of stored information during sleep helps stabilize memories, preventing forgetting of important information (Born & Wilhelm, 2012; Marr, 1971; Paller, 1997; Paller et al., 2020).

Studies using Targeted Memory Reactivation (TMR) have provided evidence for this hypothesis by demonstrating that selectively reactivating memories during sleep can strengthen them (Oudiette & Paller, 2013). In TMR experiments, learning is associated with a sensory cue, which is subsequently presented during sleep without awaking the sleeper. Cue presentation can lead to reactivation of memory content in the cortex and hippocampus (Bendor & Wilson, 2012; Cairney et al., 2018; Wang et al., 2019). After waking, reactivated memories are typically remembered better than those not reactivated, demonstrating that memory reactivation during sleep can strengthen memory, a finding that has been confirmed by meta-analysis (Hu et al., 2020).

Experiments with TMR have shown that it is a useful tool for investigating questions in memory research and potentially as an intervention for cognitive enhancement. For example, TMR can improve retention of information learned in a classroom setting (Gao et al., 2020), and facilitate learning of motor skills (Cheng et al., 2021; Johnson et al., 2019). Researchers have therefore proposed that TMR may be useful to enhance memory and to augment therapies that depend on learning, like rehabilitation (Oudiette & Paller, 2013; Paller, 2017).

A major barrier to expanding use of TMR is that the technique requires experimenters to control presentation of cues while monitoring sleep using polysomnography. In this way, cues can be presented in a particular sleep stage without arousing the participant from sleep. A specialized sleep facility and extensive training of operators is required for this online sleep scoring, and participants must sleep in an environment that differs in many ways from their typical sleeping environment at home.

These requirements impose substantial limitations on TMR experiments. For example, very few studies have examined the effects of multiple TMR sessions, owing largely to logistical difficulties. Standard TMR requirements also make it impractical to study or use TMR in clinical therapy across multiple sessions. To surmount these limitations, new ways to perform TMR in participants’ own homes are needed, ideally using an automated system that does not require direct control by an operator.

### Previous research on TMR outside of the sleep lab

Previous research on home-TMR can be divided into two categories. With unsupervised approaches, TMR cues are automatically presented during sleep irrespective of sleep stage. In supervised approaches, there is an attempt to present TMR cues in a specific sleep stage.

Unsupervised home-TMR has shown mixed results in improving cognition and memory. Ritter and colleagues (2012) found that memory reactivation during sleep could enhance creative problem solving; the researchers reactivated a problem-solving task using a plug-in scent diffuser during overnight sleep. While they slept, participants received either an odor linked to the task, an irrelevant odor, or no odor, and those who received task-linked odors produced solutions that blinded raters judged as more creative. Similarly, Neumann and colleagues (2020) found that TMR with an olfactory cue (incense sticks placed near the head while sleeping) could improve vocabulary learning in children. Other unsupervised TMR experiments, on the other hand, did not find benefits consistent with the TMR literature. Donohue and Spencer (2011) found that TMR using a continuous ocean sound played while participants slept overnight did not improve memory for word pairs. Göldi and Rasch (2019) found no effect of TMR when foreign vocabulary was cued 30 minutes after sleep onset. However, a further analysis showed that TMR benefitted memory for participants who reported that their sleep was undisturbed, but not those who reported that sound cues disturbed their sleep.

Our recent experiments using polysomnographic recordings in the lab environment substantiated the notion that Göldi and Rasch (2019) put forward—that TMR does not improve memory when sleep is disrupted by sounds. One study showed that a TMR benefit for learning face-name associations was reduced when TMR sounds disrupted sleep (Whitmore et al., 2022). Furthermore, deliberately disrupting sleep with loud sound cues reverses the TMR effect on spatial recall, selectively weakening reactivated memories (Whitmore & Paller, 2022). Therefore, we suggest that unsupervised TMR may tend to be ineffective because the intensity and timing of cues cannot be adjusted to avoid disrupting sleep.

Accordingly, supervised home TMR using a sleep sensor may be superior to unsupervised TMR. In two prior experiments with home TMR, we used a modified Zeo system (Shambroom et al., 2012) with electrolyte-filled electrodes for forehead EEG recordings used to control sound presentations. One study showed an impact of TMR on feelings of ownership and proprioceptive drift in the rubber-hand illusion (Honma et al., 2016). The other showed effects on creative problem solving (Sanders et al., 2019).

### Designing a home TMR system

Based on previous research and pilot testing, we identified key needs for a home TMR system. These included the ability to target specific sleep stages, \ robustness signal problems such as poor contact quality, and minimal reliance on proprietary or black-box technology. The system must also be comfortable, avoid disturbing sleep, and be easy for participants to use.

As no complete system currently exists meeting these requirements, we developed a new TMR system that we call SleepStim, based on consumer devices, specifically an Android phone and a smartwatch (Fitbit Versa). As sleep stages can be decoded from heart rate and wrist movement (Beattie et al., 2017; de Zambotti et al., 2018; Faust et al., 2019), we developed a custom algorithm using this information in real time to identify periods of N3 sleep. Our aim was to trigger TMR cueing during these periods, rather than to discriminate all sleep stages. We then tested whether TMR with SleepStim could improve memory for object-location associations as observed in previous TMR studies (e.g., Rudoy et al., 2009).

### Experiment 1 Methods

Figure 1 shows a diagram of the 5-day procedure. On the first day, we provided participants with a Fitbit and an Android smartphone. To allow us to correlate sleep-physiology features with behavioral results, a Dreem-2 headband (Arnal et al., 2020) was also provided. On the second day, participants learned arbitrary screen locations for 50 objects shown on a grid on the smartphone. Each object appeared with a distinct sound naturally associated with the object. On the second, third, and fourth night, sound cues for half of the objects were presented during sleep. Memory was tested in the morning of the third, fourth, and fifth day. We predicted that participants would recall locations more accurately for objects reactivated during sleep compared to those not reactivated, replicating the typical effect of TMR on spatial memory.

**Figure 1.**
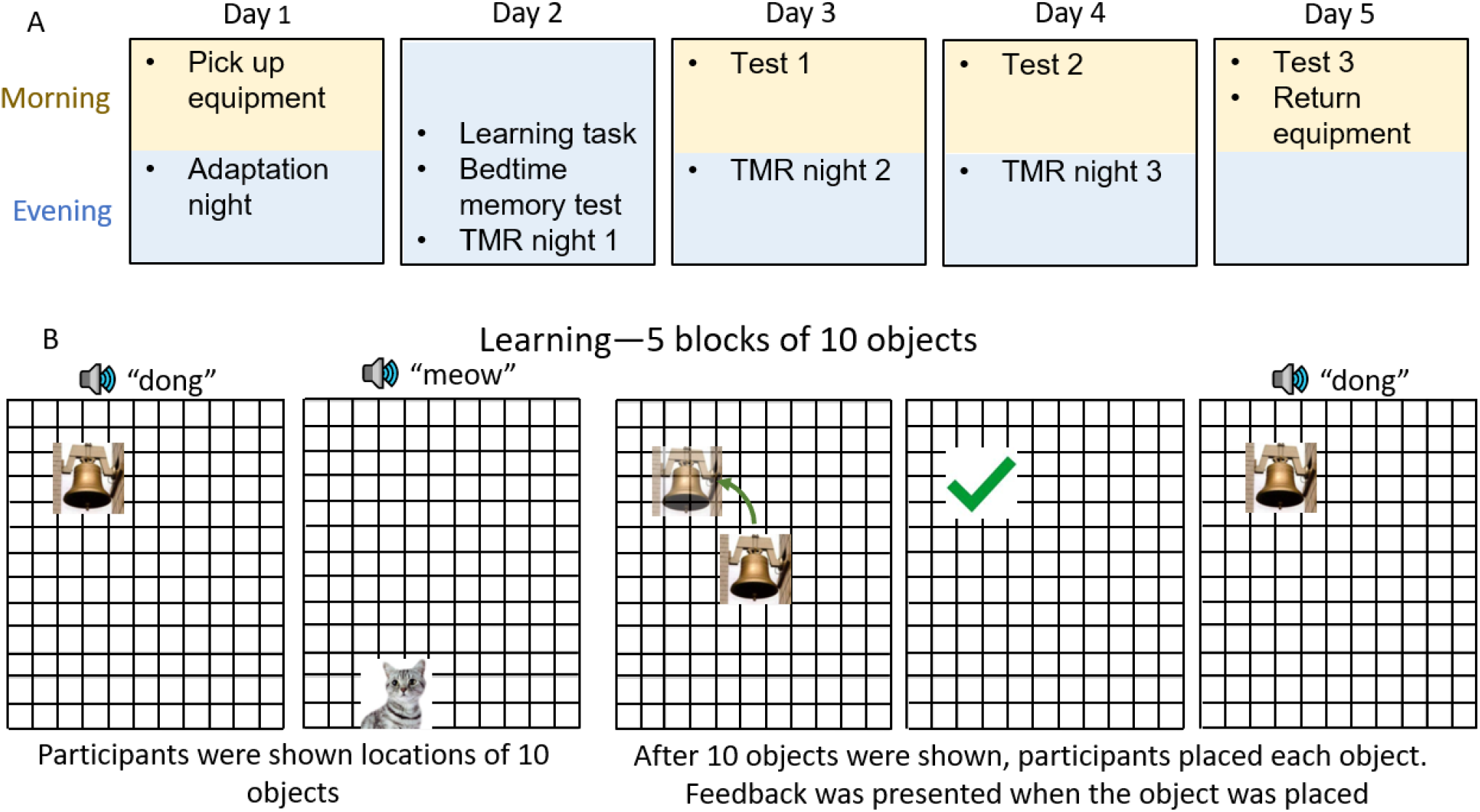
A. Sequence of events in the study. On day 2 all tasks were completed in the evening.B. Procedures for the learning phase, including presentation of objects (left) and trials of location recall with the drop-out method (right). The sound of each object was played whenever the object appeared on the screen in its target location. The memory test used the same procedure for location recall except that participants were not given feedback or shown the correct location of the objects, and the sounds were not played.

### Participants

We collected data from 120 adults who we recruited using flyers placed on campus. The protocol was approved by the Northwestern University IRB. Participants provided written informed consent and were paid for their time. Prior to data analysis, the following inclusion criteria were applied.

- Completion of training, the bedtime memory test, and at least one morning memory test
- During sleep, at least 25 cues were presented
- No more than four stimuli were presented when the Fitbit read a heart rate of zero (indicative of a poor heart rate signal)
- Objects were correctly allocated to cued and uncued conditions (which did not happen for 3 participants due to a bug in the allocation algorithm)

This process yielded data from 61 participants for analysis. Participants were between 18-25 years old (mean=20.6 years) and 28% of them were male.

### Procedure

**Day 1**. Participants picked up the equipment and were instructed on the procedure and how to use the smartphone app. They wore the Fitbit and the Dreem-2 that night to allow for acclimation to the equipment. The phone played continuous white noise overnight. Participants used a slider in the app to set the white-noise intensity to a comfortable level. This intensity setting was used as the initial setpoint for sounds played during the night. Using the algorithm described in detail below, the app controlled presentation of a control sound (electronic ding) intended to help participants adapt to the potential disruption of sound presentations. The goal was to reduce sleep disruption from experimental sounds presented on subsequent nights.Targeting slow-wave sleep, the phone repeatedly played an electronic ding sound that was unrelated to the memory task.

**Day 2**. Using the phone, participants completed the learning phase at a mean time of 10:24 PM [SEM 71 min]. In this task (described in Figure 2), a grid covered the phone’s entire screen, and participants learned the correct locations of objects on the grid. The app recorded accuracy and response times during each phase.

There were 5 blocks of trials, each with 10 objects. First, the participant was shown the correct locations of the 10 objects in that block. Then, each object appeared in the center of the screen in a random order, and the participant attempted to move it to the correct location. The participant then received feedback consisting of a red X (if incorrect) or a green checkmark (if correct) at the location where they positioned the object. The feedback was presented for 2 seconds, after which the object was displayed in the correct position for 3 seconds. The placement was considered correct if the object was placed within 120 pixels (∼2 cm) of the correct location; correct objects were dropped from the rotation. A block ended when the participant placed all objects correctly. The phone played the sound associated with each object when it first appeared on the screen, and when the correct location was shown in the feedback phase.

**Figure 2.**
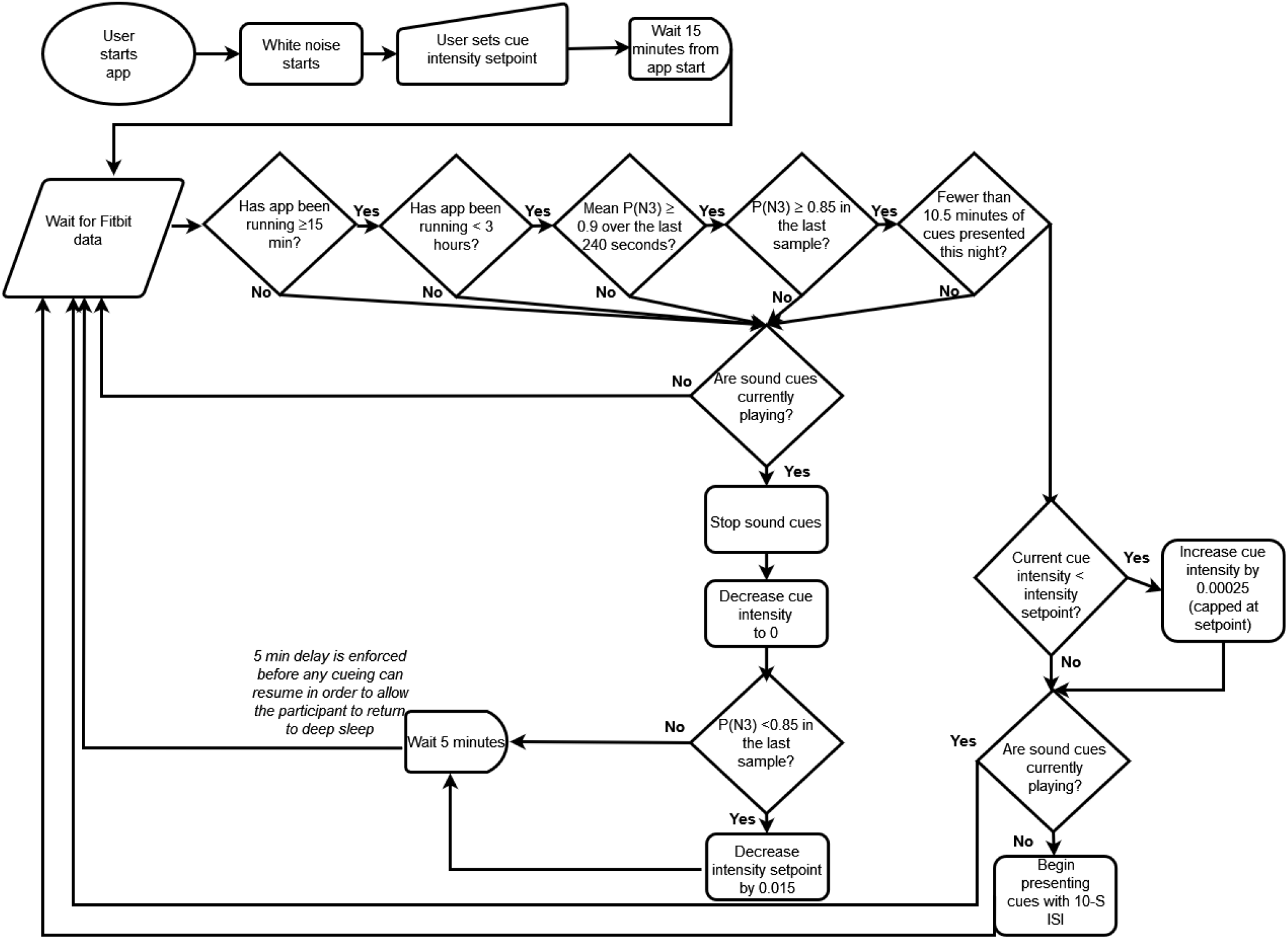
Flow chart illustrating how sounds are controlled in the SleepStim system. P(N3) is the probability that the participant is in N3 sleep as determined by the neural network classifier. The Fitbit transmits data once per second; if no Fitbit data is received for 10 seconds (indicating loss of signal), the sounds are turned off. A rapid drop in P(N3) while sounds are playing suggests the sounds aroused the participant and the intensity may be set too high, therefore the intensity setpoint is decreased if P(N3) drops below 0.85 while sounds are playing.

Participants began the bedtime memory test shortly after completing the learning (mean delay 11 min, SEM=5 min). In this test, all 50 objects were presented sequentially in the center of the screen, in random order, and the participant attempted to move each object to its correct location. Unlike in the learning phase, no feedback was given.

After participants completed their bedtime memory test, the app selected 25 objects to be cued during sleep using a matching algorithm to minimize the difference in bedtime memory performance between two sets of objects (to be cued and uncued). Objects were sorted by memory error and then assigned in alternating order (i.e., 1=cued, 2=uncued…). Because some differences remained after this assignment, the app also counterbalanced participants so that the assignment procedure started with cued in half of the participants and uncued in the other half. As expected, there was no difference in recall accuracy between cued and uncued objects in the bedtime memory test [Wilcoxon signed-rank test, z(60)=0.8,p=0.42].

Shortly before going to sleep, the participant put on the Fitbit and Dreem-2, started the TMR app, and calibrated white-noise intensity. During the night, and on all subsequent nights, sounds linked to the 25 objects in the cued condition were presented during sleep.

**Days 3 and 4**. Participants completed a memory test in the morning. The test was identical to the memory test on Day 2, except with a different random order of objects. During sleep, they used the Fitbit, Dreem-2, and TMR app as on previous nights.

**Day 5**. Participants completed a final memory test in the morning and returned the equipment. When returning the equipment, we asked participants whether they remembered hearing any of the sounds from the memory task while they were sleeping.

### Automated TMR

Sounds were played at 10-second intervals when N3 sleep was detected. Detection was operationalized as (a) a high value for the probability of N3 averaged over the most recent 240 s, P(N3) ≥ 0.9, and (b) the most recent value for P(N3) ≥ 0.85. Cue start/stop and cue sound intensity was controlled by the algorithm shown in Figure 2.

SleepStim was limited to presenting sounds when P(N3) was high within a time interval from 15 minutes to 3 hours after the time the system was turned on. The system was also limited to a maximum time of stimulation of 10.5 minutes. These constraints were imposed to minimize the chance of disrupting sleep with the sounds, and are consistent with the protocols used in previous lab-TMR studies (Rudoy et al., 2009).

### Memory performance measurement

We measured *memory change* as the ratio of mean spatial error at a morning memory test to mean spatial error at the bedtime memory test (e.g., mean test1 error / mean bedtime test error). We computed this statistic separately for cued and uncued objects.

For each test, we computed the *TMR effect* as the memory change for cued objects minus the memory change for uncued objects. A negative value indicates a benefit of TMR for memory. For example, a TMR effect of −0.1 implies that the increase in error for cued objects was 10% lower than the increase in error for uncued objects. We determined whether TMR effects differed significantly from zero using the Wilcoxon signed-rank test, a non-parametric one-sample test.

In the primary analysis of the TMR effect, we examined memory performance on the last test taken. Whereas participants were asked to take three memory tests, some failed to do so on one or more mornings. Therefore, the primary analysis focused on the memory error from the last test, and we treated the number of prior memory tests and the number of nights cued as predictors in our model.

In the secondary analysis, we considered only participants who completed all three memory tests (*n* = 41). We evaluated the effect of TMR at each test to observe how it changed over time.

### Controlling for effects of initial memory performance

We observed that TMR effects were correlated with the pre-sleep difference in memory performance between cued and uncued objects (Figure 3), which could be interpreted as regression to the mean. That is, the larger the cued/uncued difference initially, the more likely this difference is reduced on the subsequent test. Because this effect adds variability that could obscure other correlations, we controlled for this effect before analyzing relationships between the TMR effect and other variables. In this procedure, we used linear regression to isolate the relation between the TMR effect and initial memory performance differences between cued and uncued objects, computed separately for each test. The residual effect after covarying out this relationship was calculated as the *corrected TMR effect*.

**Figure 3.**
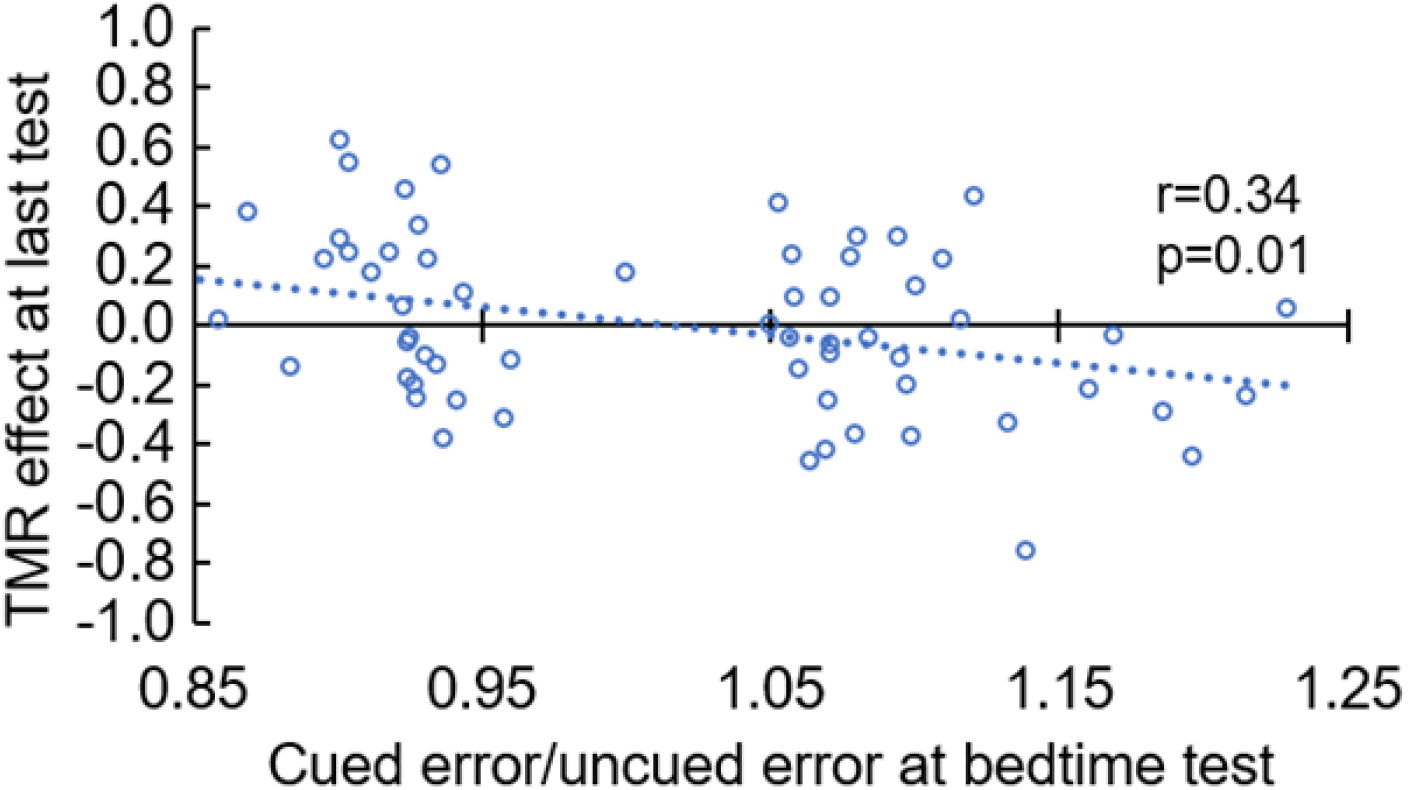
An example of the linear regression used to control for variation in memory performance in the bedtime test in Experiment 1. The bimodal distribution of bedtime test scores resulted from the procedure used to assign objects to cued and uncued conditions.

### Dreem-2 sleep staging

We used data from the Dreem-2 headset to compute the time participants spent in each sleep stage, as well as the percentage of cues delivered in each sleep stage. Some participants did not have sufficient high-quality data for staging (by the proprietary Dreem algorithm), so only a subset of 45 participants were included in these analyses.

### SleepStim system

A custom application running on the Fitbit acquired data once per second. The data consisted of heart rate in beats per minute, acceleration on X, Y, and Z axes, and rotation on these axes from the accelerometer and gyro, respectively. The data were transmitted via Bluetooth to the paired phone. The first step of processing on the phone was feature extraction, as schematized in Figure 4. Briefly, the phone computed a time-frequency representation of the last 240 seconds of accelerometer, gyro, and heart-rate data. The result was a time-frequency matrix, quantifying variability as a function of both time (number of prior seconds) and frequency. This transformation is similar to that used in other sleep-staging algorithms (Beattie et al., 2017), and is useful because it allows for characterization of various sleep phenomena (e.g., high-frequency vs. low-frequency heart rate variability). Because time-frequency variability was highly correlated on all axes, only the Z axis of the accelerometer and gyro signals was used in computing the time-frequency representation. Following feature extraction, the time-frequency features along with current values from all sensors and total motion integrated over the last 240 seconds were input to an artificial neural network classifier trained to predict the probability of N3 sleep. For each sample (each second), the network produced a value, P(N3), corresponding to the probability of N3 sleep.

**Figure 4.**
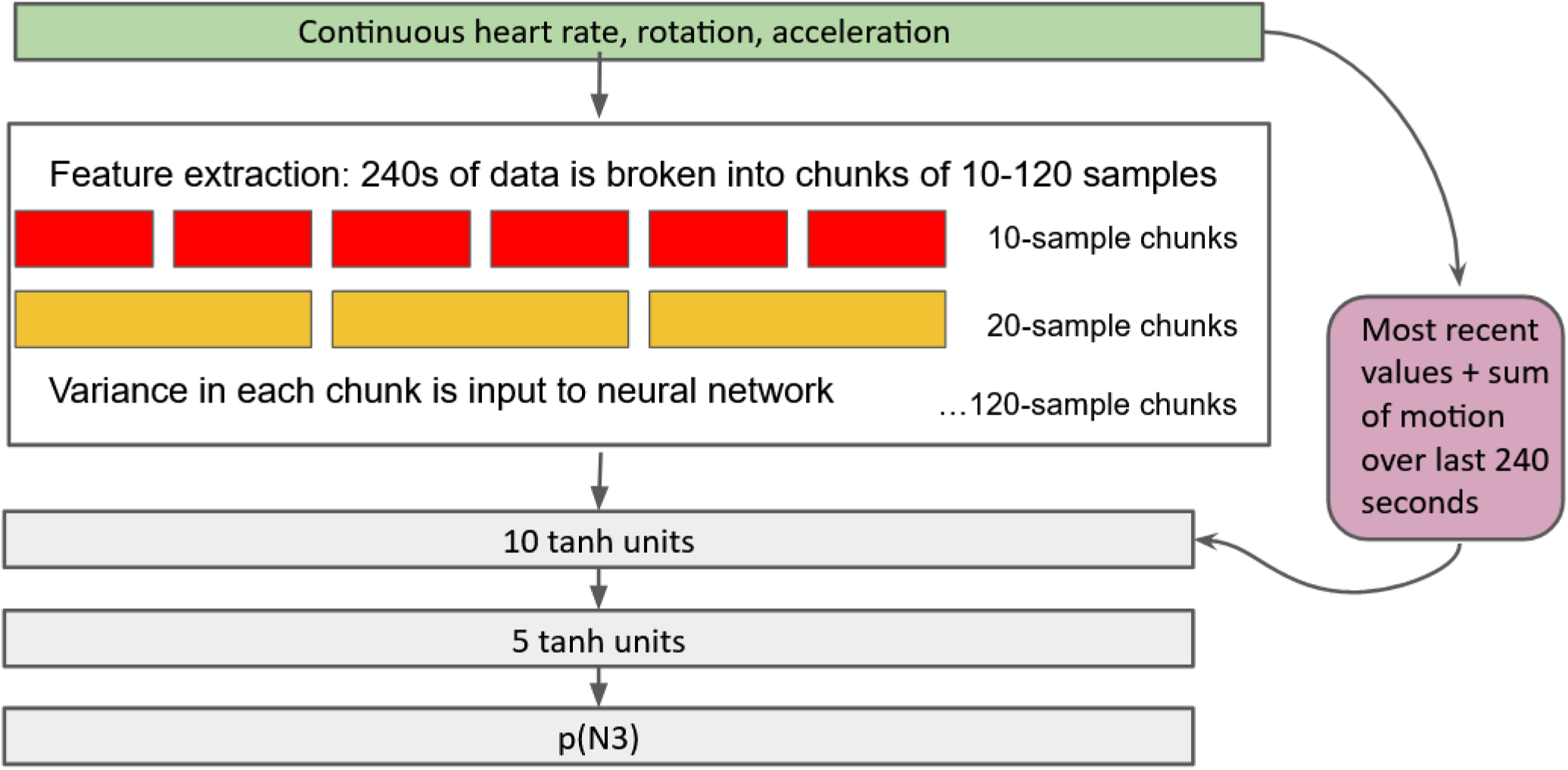
Schematic of the feature-extraction system and neural network. Variance was calculated using the standard deviation of each chunk.

### Neural network training and testing

We trained the neural network on a dataset for 24 participants that included Fitbit data and sleep scores from an overnight session. Half of the participants were young adults who slept in the lab overnight for an unrelated study and half were middle-aged adults who slept at home. For the young adults, sleep stages were determined by manual scoring of polysomnographic data; for middle-aged adults sleep stages were determined using the automatic scoring built into the Dreem-2.

Prior to training, we computed features for the Fitbit data as described above. To speed training, we subsampled the data by a factor of 5, to yield one sample every 5 seconds. Preliminary testing showed that this did not meaningfully affect classifier accuracy, likely because successive samples contained mostly redundant information. In total, 178,948 observations were included in the dataset.

We then trained a perceptron neural network classifier with two hidden layers to predict whether each second would be scored as N3 based on the Fitbit features. Training was performed using the Neural module of JMP 15.2.1 (2019) using the “squared” regularization penalty. To evaluate the network’s overall performance in classifying N3 sleep, we also trained a separate version of the model with one-third of the subjects (50,425 observations) held out from training as a validation set. The model achieved an area under the curve of 0.77 in classifying sleep as N3 or non-N3, indicating that it exceeded chance performance. We also evaluated alternative classifier schemes, including linear discriminant analysis and a convolutional neural network. Of these, the two-layer perceptron combined with our feature extraction algorithm performed the best.

## Experiment 1 Results

### TMR cues were effectively targeted to N3 sleep

For 45 participants with EEG recordings of sufficient quality to permit sleep staging during cueing, we compared the percent of cues delivered in each sleep stage to the percent of overall time spent in that sleep stage. This analysis served as an independent test that the algorithm targeted N3 sleep in a new group of participants following the original test and validation set.

Results shown in Figure 5 revealed that SleepStim successfully targeted N3. Compared to the total time in each stage, the time when cues were played was more likely to be N3 [t(44)=3.56, p<0.001] and less likely to be classified as N2 [t(44)=2.26, p=0.03] or REM [t(44)=2.61, p=0.01]. Although N2 was underrepresented in the cued sleep, a substantial number of cues were presented in N2 due to the higher base rate of N2 sleep. We did not observe differences between total sleep and cued sleep in wake or N1, which may be because these stages were rarely observed in the training set, providing little opportunity for the model to learn how to identify them.

**Figure 5.**
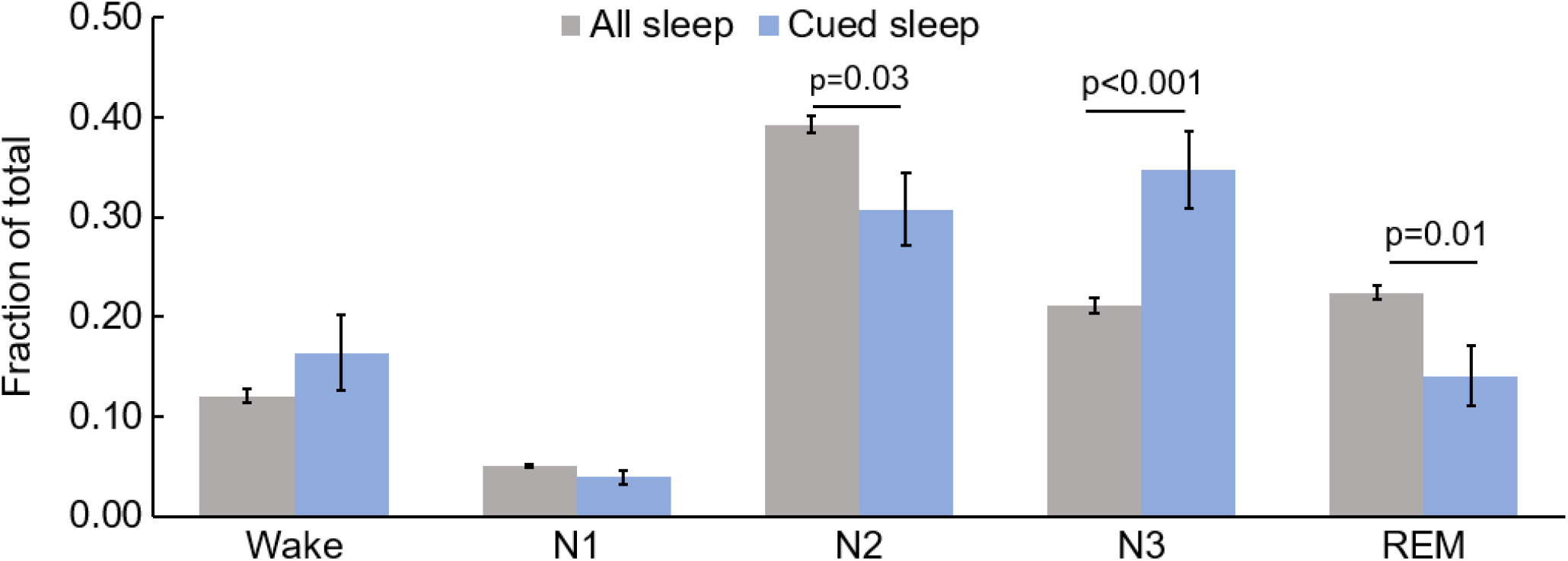
Results showing that the SleepStim system predominantly delivered cues during N3 sleep. Gray bars show the proportion of time spent in each sleep stage and blue bars show the distribution of sleep stages when cues were delivered. Whereas N3 comprised 21.1% of sleep, 34.7% of the cues were delivered in this stage, and 65.5% of the cues were delivered in stages N2 or N3.

### Participants generally did not notice TMR cues

In the full sample (including participants who did not pass inclusion criteria), 16/120 participants (13%) reported hearing at least one sound from the memory task. No participants reported that the sounds disrupted their sleep or woke them. Among the participants included in analysis, 7/61 (11%) reported hearing at least one sound.

### Participants efficiently learned and retained object locations

Participants required a mean of 1.61 [SEM=0.09] repetitions per object in the learning phase to reach criterion. In the bedtime test, participants’ mean accuracy surpassed the criterion (Figure 6), indicating that the learning procedure created an effective memory at a short delay.

**Figure 6.**
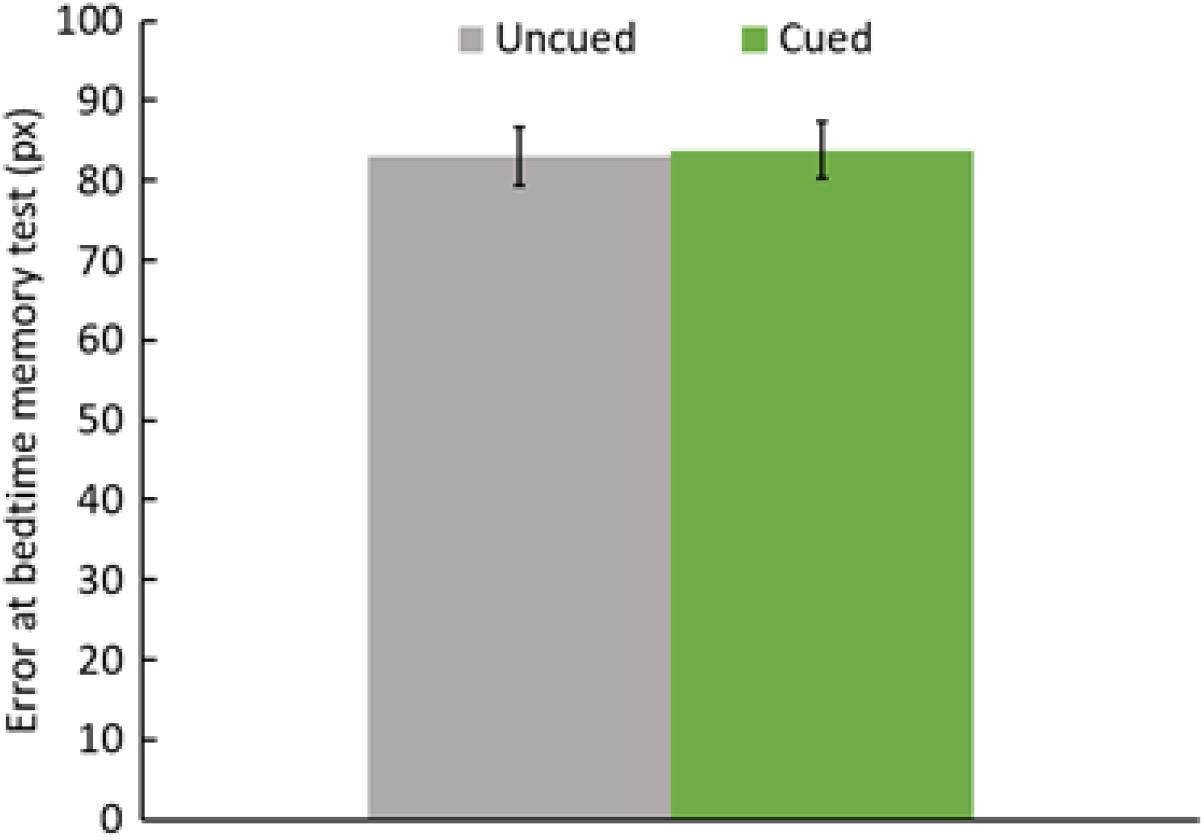
Error at the bedtime test remained below the learning criterion (120 pixels). Error did not differ between cued and uncued conditions [Wilcoxon signed-rank test; mean difference=0.76 pixels, z(60)=0.8,p=0.42]

### Recall accuracy declined but seemed uninfluenced by TMR

Mean spatial error increased from 83 pixels at bedtime test to 105 pixels at last test, indicating significant forgetting [*t*(60)=9.48, *p*<0.001]. No significant TMR effect was found at the last test or at any of the individual time points (Figure 7).

**Figure 7.**
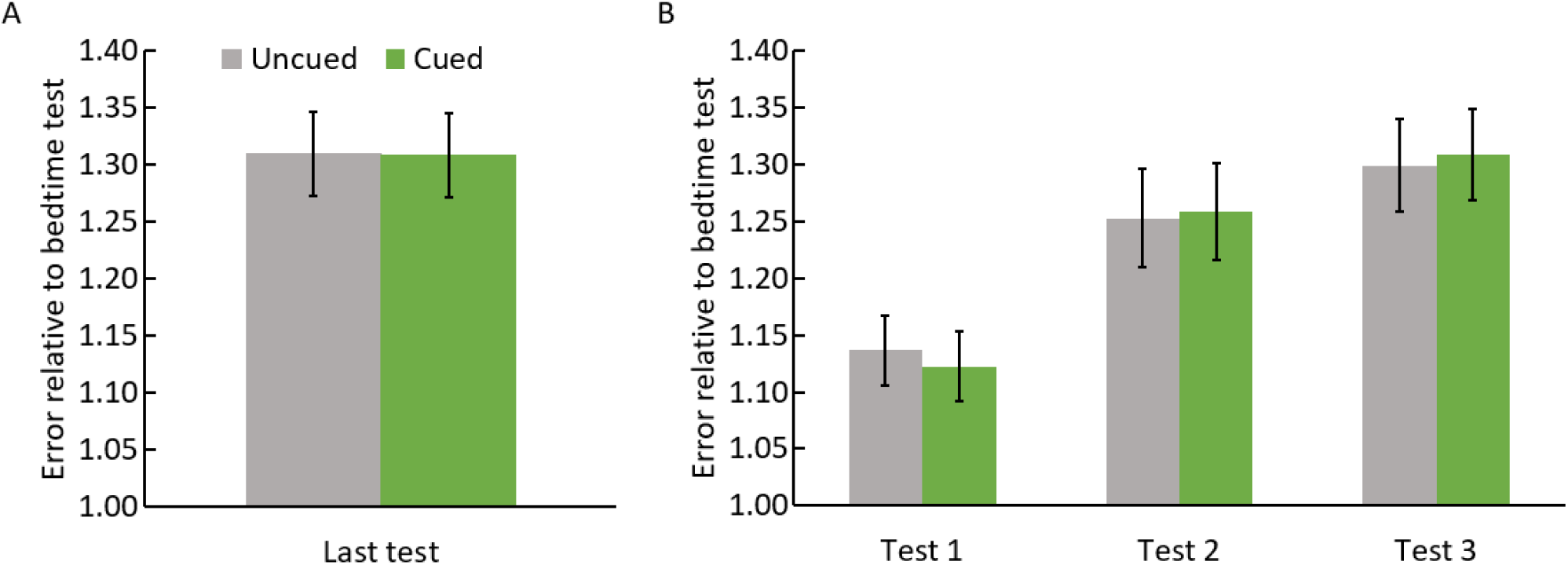
Mean spatial error increased by about 30% for both cued and uncued objects at the last test (compared to the bedtime test immediately after learning). No significant difference in error was found between cued and uncued objects. B. In participants who completed all 3 morning tests (*n*=41), error continued to increase throughout the experiment, reflecting forgetting. Error bars reflect the SEM for the within-subjects analysis of cued error-uncued error.

### TMR effect was associated with cue-sound intensity and sleep-stage targeting

One factor shown to impact TMR effects in prior studies is level of sleep disruption (Whitmore et al., 2022). Therefore, we hypothesized that sleep disruption caused by cues might have influenced the extent to which participants responded here. One potentially relevant parameter is sound intensity. Given that sleep could be disrupted when sounds were too loud, we quantified the maximum intensity used overnight. The memory benefit from TMR was significantly correlated with maximum cue intensity and marginally correlated with the percentage of cues delivered in stage N3 (Figure 8, Table 1).

**Table 1.**
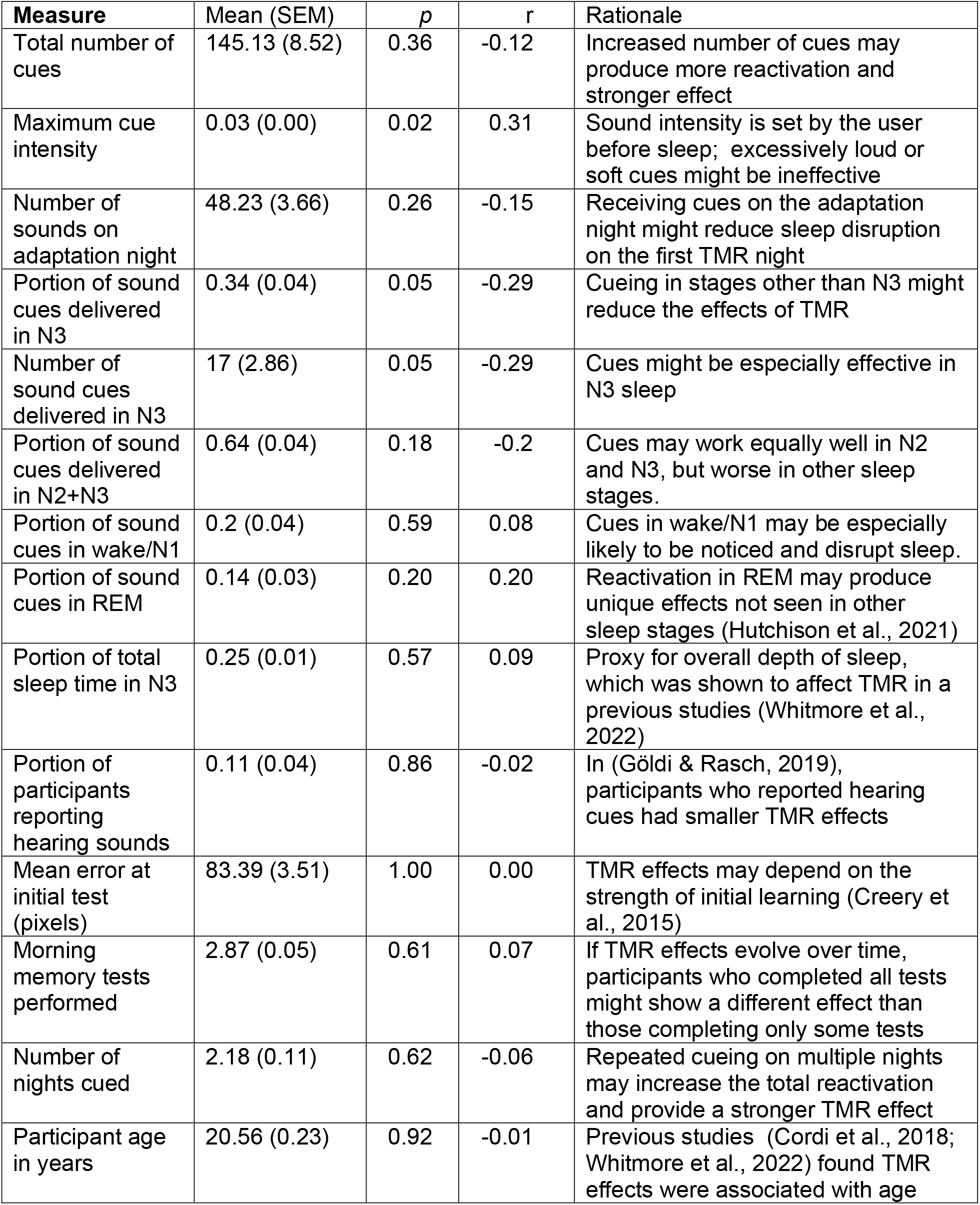
Correlations between the corrected TMR effect and sleep/participant variables. Correlation is a linear regression. Sign of the r value indicates the direction of the correlation; a negative r indicates higher values of the independent variable are associated with more benefits of TMR for memory

| Measure | Mean (SEM) | *p* | *r* | Rationale |
| --- | --- | --- | --- | --- |
| Total number of cues | 145.13 (8.52) | 0.36 | -0.12 | Increased number of cues may produce more reactivation and stronger effect |
| Maximum cue intensity | 0.03 (0.00) | 0.02 | 0.31 | Sound intensity is set by the user before sleep; excessively loud or soft cues might be ineffective |
| Number of sounds on adaptation night | 48.23 (3.66) | 0.26 | -0.15 | Receiving cues on the adaptation night might reduce sleep disruption on the first TMR night |
| Portion of sound cues delivered in N3 | 0.34 (0.04) | 0.05 | -0.29 | Cueing in stages other than N3 might reduce the effects of TMR |
| Number of sound cues delivered in N3 | 17 (2.86) | 0.05 | -0.29 | Cues might be especially effective in N3 sleep |
| Portion of sound cues delivered in N2+N3 | 0.64 (0.04) | 0.18 | -0.2 | Cues may work equally well in N2 and N3, but worse in other sleep stages. |
| Portion of sound cues in wake/N1 | 0.2 (0.04) | 0.59 | 0.08 | Cues in wake/N1 may be especially likely to be noticed and disrupt sleep. |
| Portion of sound cues in REM | 0.14 (0.03) | 0.20 | 0.20 | Reactivation in REM may produce unique effects not seen in other sleep stages (Hutchison et al., 2021) |
| Portion of total sleep time in N3 | 0.25 (0.01) | 0.57 | 0.09 | Proxy for overall depth of sleep, which was shown to affect TMR in a previous studies (Whitmore et al., 2022) |
| Portion of participants reporting hearing sounds | 0.11 (0.04) | 0.86 | -0.02 | In (Göldi & Rasch, 2019), participants who reported hearing cues had smaller TMR effects |
| Mean error at initial test (pixels) | 83.39 (3.51) | 1.00 | 0.00 | TMR effects may depend on the strength of initial learning (Creery et al., 2015) |
| Morning memory tests performed | 2.87 (0.05) | 0.61 | 0.07 | If TMR effects evolve over time, participants who completed all tests might show a different effect than those completing only some tests |
| Number of nights cued | 2.18 (0.11) | 0.62 | -0.06 | Repeated cueing on multiple nights may increase the total reactivation and provide a stronger TMR effect |
| Participant age in years | 20.56 (0.23) | 0.92 | -0.01 | Previous studies (Cordi et al., 2018; Whitmore et al., 2022) found TMR effects were associated with age |

**Figure 8.**
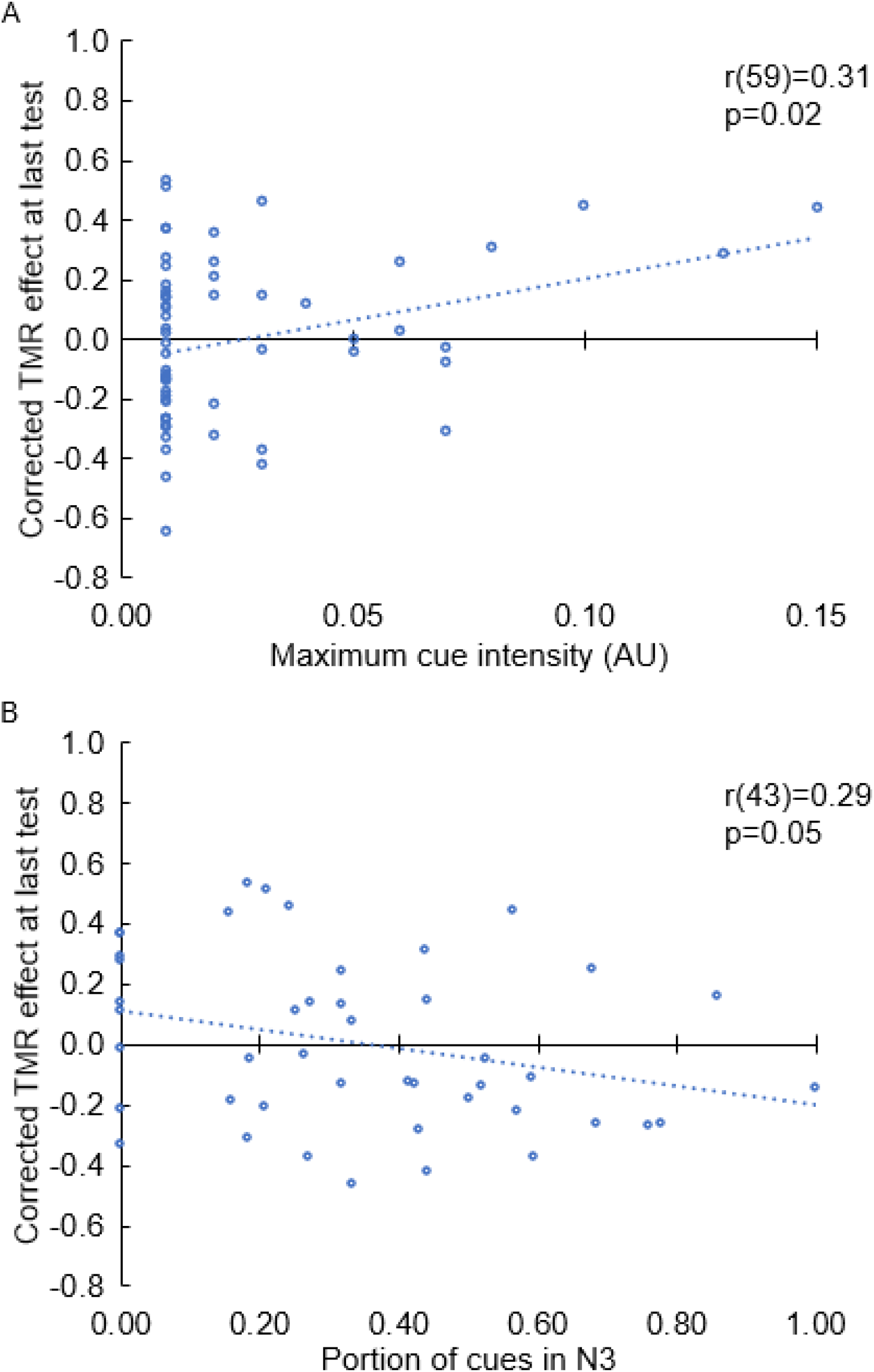
A. Correlation between corrected TMR effect and maximum cue intensity. B. Correlation between corrected TMR effect and proportion of cues in N3.

### Comparing TMR with optimal versus non-optimal parameters

Given the correlational results, we explored individual differences further by considering whether TMR might have a larger benefit in participants cued with optimal parameters, defined as receiving at least 25 sound cues on the adaptation night and using a relatively low maximum sound intensity (<0.02). We opted to select these participants because these two factors, adaptation procedures and sound intensity, could be directly controlled by the experimenter to reduce sleep disruption. Differences between the two groups were nonsignificant (Figure 9), but we did observe trends where the TMR effect was larger for the optimal-cued participants at last test [Mann-Whitney *U* test, U(60)=356, *p*=0.12] and at test 3 [U(52)=258, *p*=0.11]. Neither the optimal or non-optimal group showed a significant effect of TMR at the last test or at test 3.

**Figure 9.**
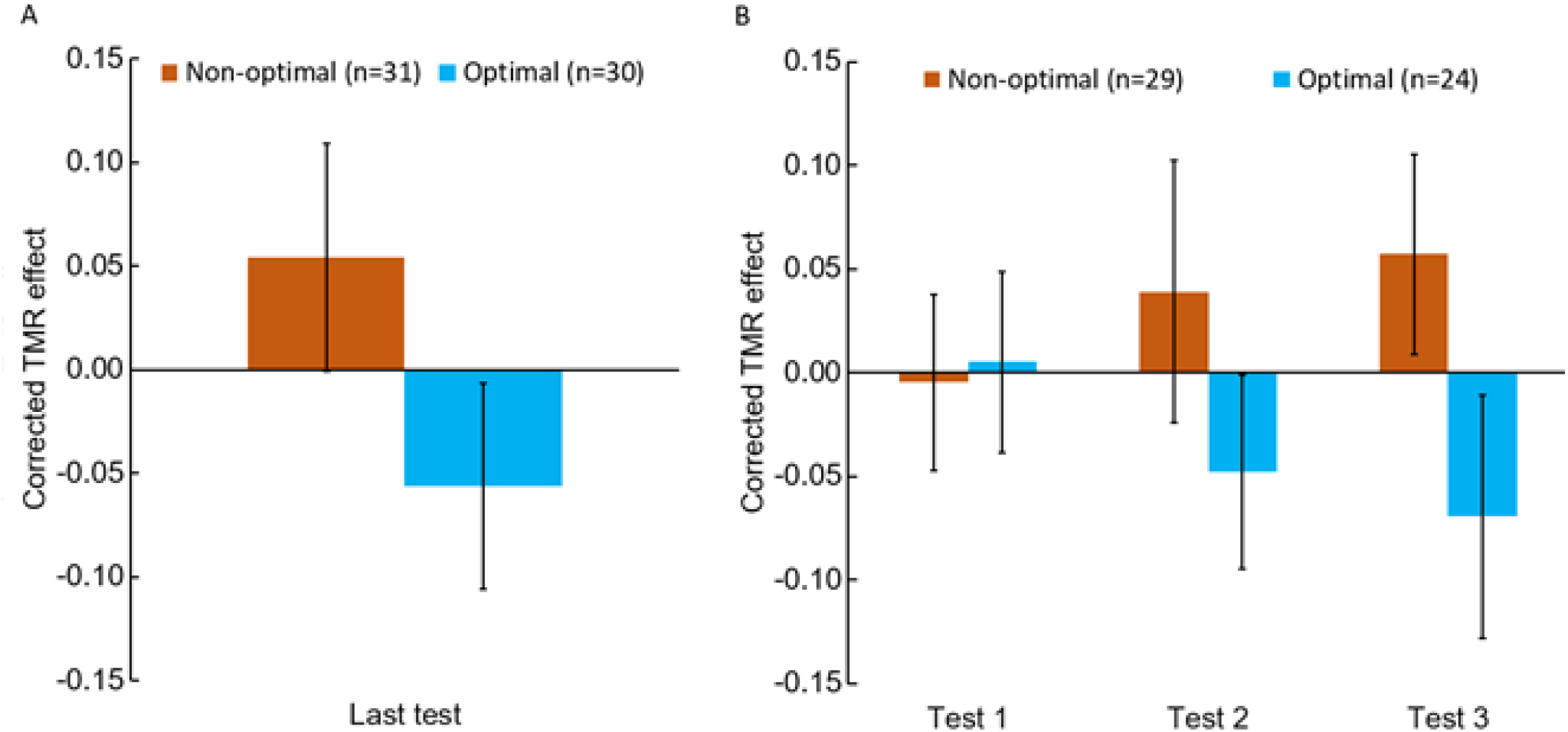
Participants cued with optimal parameters showed a trend towards a benefit from TMR. Participants cued with non-optimal parameters showed a trend towards a detrimental effect of TMR. Results are shown for participants on their last test (A) and for all 3 tests (B).

### Experiment 2 Methods

Because our initial experiment suggested that SleepStim could improve memory contingent on low cue intensity, we conducted a follow-up study implementing an improved method. This experiment was identical to the original experiment, except for the following modifications.

- Participants could not set initial intensity higher than 0.02 A new algorithm required participants to receive at least 25 adaptation cues before they could begin the memory test, and if 25 cues were not presented, the adaptation procedure was administered again
- A new algorithm required participants to receive at least 25 adaptation cues before they could begin the memory test, and if 25 cues were not presented, the adaptation procedure was administered again
- Participants could receive up to 30 minutes of cueing per night (compared to 10.5 minutes in Experiment 1)
- We improved the algorithm for allocating objects to cued and uncued conditions, which matched conditions more closely, obviating the need for controlling for pre-sleep memory performance in the analysis

### Participants

Participants were recruited and paid using the same methods as the prior experiment. We collected data from 44 participants, and of these, 24 passed inclusion criteria and their data were included in analysis. Participants ranged from 18-31 years old (mean=21.6 years) and 29% were male.

## Experiment 2 Results

*Participants rarely reported hearing sounds*

In the full sample, 3/44 participants (7%) reported hearing TMR sounds during sleep. Among participants included in analysis, 2/24 (8%) reported perceiving TMR sounds, and 1 reported that the sounds disturbed their sleep.

*TMR improved spatial memory at last test*

As shown in Figure 10, the improved TMR protocol significantly improved memory for cued objects relative to uncued objects at the last test [Wilcoxon signed-rank test; z(23)=-2.69, p=0.007]. For participants who took all three memory tests (*n*=18), a significant difference between cued and uncued conditions emerged at the second memory test and persisted in the third test [Wilcoxon signed-rank test; z(17)=-1.63, −2.24,−2.29, *p*=0.103,0.025,0.022 for test 1, 2, and 3, respectively].

**Figure 10.**
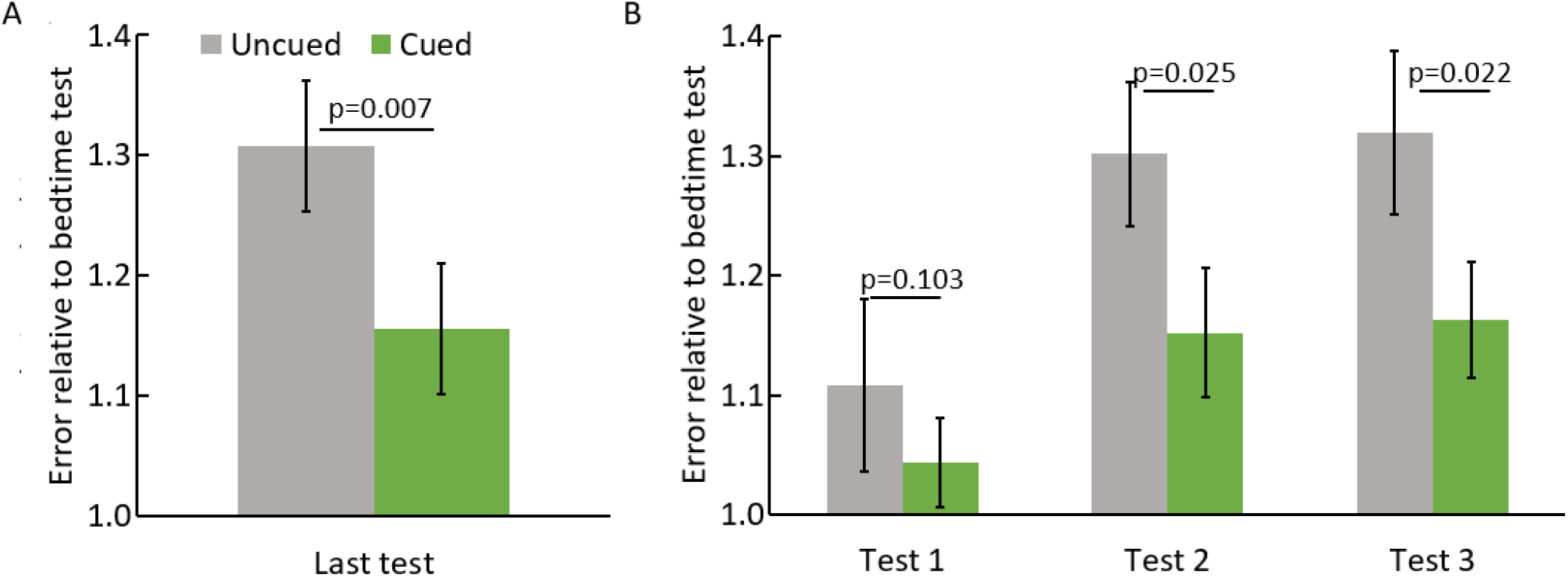
A. TMR effects at the final memory test. B. TMR effects at each time point, for the participants who performed all three morning memory tests. Error bars reflect the SEM for the within-subjects analysis of cued error versus uncued error. Figure S1 shows data for individual participants.

## Discussion

Given that studies in the home environment would greatly expand research and applications related to memory processing during sleep, we designed and tested SleepStim, a novel wearable system for presenting auditory cues during sleep. Our results confirmed that SleepStim can target deep sleep and produce memory benefits that mirror those achieved via memory reactivation in sleep labs equipped with polysomnographic equipment. We used a wearable device for obtaining EEG data from the forehead to validate our procedure, but the TMR method we devised can be applied with this system using only a wrist-worn device and a smartphone, making it easy to use, efficient, relatively inexpensive, and well tolerated by most individuals. The results demonstrate that home sleep interventions with the SleepStim system are feasible and effective, provided consideration is given with respect to cue intensity and control algorithms.

An important finding in our experiment was that participants remained unaware that TMR cues were presented in almost all cases, with 13% reporting hearing the cues in Experiment 1 and 7% reporting cues in Experiment 2. This is a substantial improvement over unsupervised home TMR in past studies in our lab and others, where participants frequently reported hearing cues and having their sleep disturbed (e.g., Göldi & Rasch, 2019). Presenting cues without participants noticing is important for usability and to avoid accidentally unblinding participants in experiments where they are assigned to different conditions. This result also confirms that SleepStim can target states where participants are soundly asleep.

The ability to target deep sleep was also reflected by analysis of the times of cue delivery in relation to the automatic sleep staging provided by Dreem-2. Cues were delivered disproportionately in N3 sleep, and most of the cues not delivered in N3 were delivered in N2. In a recent meta-analysis of the TMR literature, memory benefits were found for both N2 and N3 sleep (Hu et al., 2020). In TMR experiments aimed at enhancing memory in the sleep lab environment, cues are typically presented in either N3 or a combination of N2 and N3, and memory-related sleep features like spindles and slow waves occur in both of these stages (Dijk et al., 1993). Our results show that SleepStim can target deep non-REM sleep and deliver cues without waking participants, both important advances for sleep-intervention studies in the home. Our findings also showed that better targeting of N3 was associated with stronger benefits of TMR for memory (Figure 8B), further emphasizing the importance of targeting cues to N3.

We also demonstrated that the TMR procedure at home can yield the typical effect observed in the lab, where TMR with quiet sounds improves performance in a spatial memory task (Antony et al., 2018; Creery et al., 2015; Rudoy et al., 2009; Schechtman et al., 2021; Vargas et al., 2019). We also found that loud cues reversed the TMR effect, consistent with our prior findings in a study of face-name learning (Whitmore & Paller, 2022). Accordingly, our findings suggest that home-TMR can be useful for investigating memory and perhaps in clinical applications as well. In particular, TMR at home may open up possibilities for clinical research with TMR, studies of performance enhancement over multiple nights, and TMR studies with larger numbers of participants and greater efficiency.

Our results also highlight the critical importance of auditory cue intensity and sleep disruption in home TMR. Initially we allowed users to adjust sound intensity individually; however, forcing a consistent low level, as in Experiment 2, produced larger TMR effects and presumably produces less sleep disruption. Therefore, our recommendation is that home-TMR experiments use cues barely audible in a quiet room. Optimizing methods for calibrating intensity and making adjustments during the night is an important challenge for future research.

In this research we acquired a large amount of combined data from wearable devices on the wrist and head. Such data could be useful, especially given large participant samples, for studying how sleep parameters and experimental-design factors like the number and timing of cues relate to TMR response. Likewise, improved algorithms could possibly be developed for sleep staging using wearable data.

SleepStim offers a powerful platform for future sleep research. The ability to run TMR experiments at scale outside the sleep lab can enable new fundamental and clinical studies. The ability to deliver closed-loop interventions in sleep using SleepStim may also be useful for applications beyond TMR, such as influencing dream content (Konkoly et al., 2021) or non-phase-locked entrainment to increase slow wave and spindle activity (Antony & Paller, 2016; Simor et al., 2018).

## Acknowledgments

We thank Kristin Sanders, Kara Dastrup, and Carmen Westerberg for contributing data used to train the model. Marc Slutzky, Prashanth Prakash, Vamshi Muvvala, and Soheil Borhani provided valuable input in developing and testing the approach. Funding was provided from NSF BCS-1921678, NIH/NINDS R01NS112942, NIH/NINDS T32 NS047987, and NIH/NIMH T32 MH067564.

## Supplementary information

**Figure S1.**
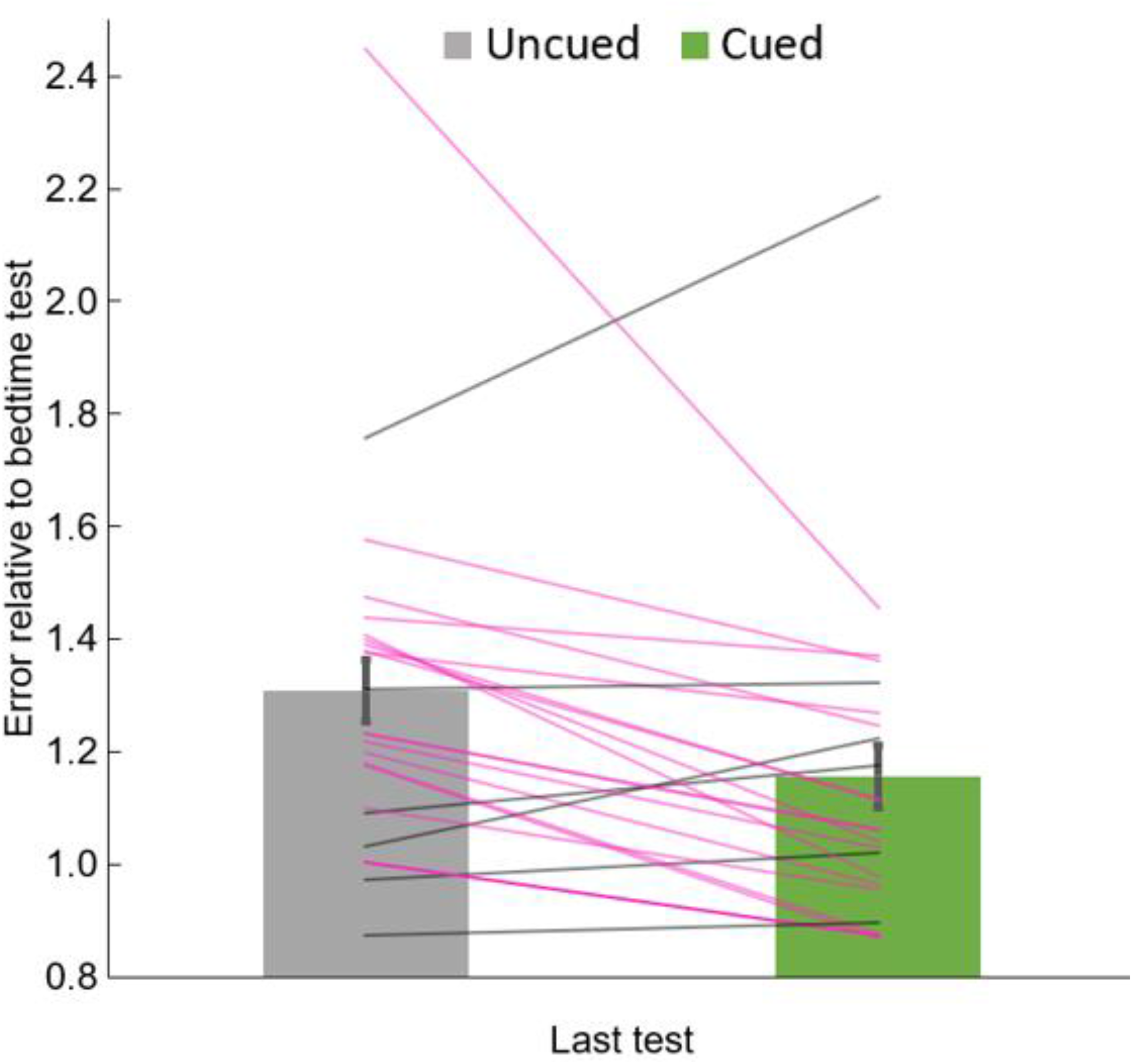
Data from Figure 10A with individual participant values. Each line represents one participant. Participants with improved memory for cued objects relative to uncued objects are shown in pink, others are shown in gray.

